# Focal Non-invasive Deep-brain Stimulation with Temporal Interference for the Suppression of Epileptic Biomarkers

**DOI:** 10.1101/2022.03.29.486252

**Authors:** Emma Acerbo, Aude Jegou, Charlotte Luff, Patrycja Dzialecka, Boris Botzanowski, Florian Missey, Ibrahima Ngom, Stanislas Lagarde, Fabrice Bartolomei, Antonino Cassara, Esra Neufeld, Viktor Jirsa, Romain Carron, Nir Grossman, Adam Williamson

**Affiliations:** Aix Marseille Univ, INSERM, Institut de Neurosciences des Systèmes, Marseille, France; Department of Brain Sciences, Imperial College London, London, United Kingdom; APHM, Timone Hospital, Department of Epileptology, Marseille, France; Foundation for Research on Information Technologies in Society, Zurich, Switzerland; Department of Functional and Stereotactic Neurosurgery, Timone University Hospital, Marseille, France; Center for Bioelectronic Medicine, Department of Medicine, Solna, Karolinska Institute, Stockholm, Sweden

## Abstract

Neurostimulation applied from deep brain stimulation (DBS) electrodes is an effective therapeutic intervention in patients suffering from intractable drug-resistant epilepsy when resective surgery is contraindicated or failed. Inhibitory DBS to suppress seizures and associated epileptogenic biomarkers could be performed with high-frequency stimulation (HFS), typically between 100 –165Hz, to various deep-seated targets such as for instance the Mesio-temporal lobe (MTL) which leads to changes in brain rhythms, specifically in the hippocampus. The most prominent alterations concern high-frequency oscillations (HFOs), namely increase in ripples, a reduction in pathological Fast Ripples (FRs), and a decrease in pathological interictal epileptiform discharges (IEDs). In the current study, we use Temporal Interference stimulation to provide a non-invasive focal DBS (130 Hz) of the MTL, specifically the hippocampus, which increases physiological ripples, and decreases the number of FRs and IEDs in a mouse model of epilepsy. Similarly, we show the inability of 130 Hz transcranial current stimulation (TCS) to achieve similar results. The method could potentially revolutionize how DBS, certainly in epilepsy, is performed, and we therefore further demonstrate the translatability to human subjects via measurements of the TI stimulation vs TCS in human cadavers. Results show the better penetration of TI fields into the human hippocampus as compared with TCS. Finally, we provide evidence of the efficacy of the specific form of Pulse-width Modulated TI (PWM-TI), implemented with square waves, which is used in this study.

**One Sentence Summary:** A non-invasive deep brain stimulation applied via temporal interference achieves the suppression of biomarkers of epilepsy in mice and is scaled to humans.

## INTRODUCTION

Epilepsy is a severe and highly prevalent neurological disease, affecting about 1% of the population worldwide *(1)*. Despite the continuous development of new antiseizure medications *(2)*, pharmacoresistance remains a major issue for about one-third of patients, who may benefit from epilepsy surgery when epilepsy is focal *(3)*. For patients with drug-resistant epilepsy not eligible for surgery or with failure of resective surgery, neuromodulation, *i*.*e*. electrical stimulation, be it invasive or not, provides an alternative option. Among neuromodulation techniques, deep brain electrical stimulation (DBS) has been increasingly explored and included in clinical practice *(4–6)*. The most common methods of DBS use implanted leads targeting thalamus, hippocampus, or other parts of basal ganglia depending on the indication and types of epilepsies *(7, 8)*. These approaches proved to be efficient, effective and well tolerated *(9, 10)*, but remain invasive and are not deprived of possible complications (*e*.*g*. neurological deficit, hematoma, infection) *(11)*. As a result, non-invasive neurostimulation techniques such as transcranial current stimulation (TCS) provide interesting perspectives and are currently on trial as therapeutic approaches in epilepsy *(12)*. However, the limited spatial accuracy and the low penetration of deepest brain regions are limitations of current TCS protocols *(13, 14)*. To address this, we utilize the method of temporal interference (TI) stimulation, a focal stimulation method which has the ability to stimulate deep brain structures with interferences at points located at significant distance from the surface electrode *(15)*.

Ideal parameters for neuromodulation regimes specifically treating epilepsy using electrical stimulation of the peripheral *(16)* or central nervous system (CNS) *(17)* have been evaluated with both animal and clinical studies *(18)*. The aim of CNS stimulation in epilepsy is to target a key region in the brain and apply electrical stimulation, traditionally with a DBS electrode, to suppress seizures and the associated epileptogenic biomarkers *(10)*. Inhibitory DBS is delivered at high-frequency stimulation (HFS), typically between 100–165Hz *(19)*. The main DBS study for seizure suppression in epilepsy was the SANTE trial, with more than 100 patients enrolled. It aimed to assess the safety efficacy of the stimulation of the anterior nucleus of the thalamus (ANT). The trial has shown HFS at 145Hz could significantly decrease seizure frequency and improve patients’ quality of life *(20, 21)*. Originally HFS at 130Hz was a parameter used in the treatment of essential tremor and Parkinson’s disease, however it has also been successfully applied in epilepsy *(22, 23)*. The same frequency has now been used effectively in several different regions of the brain, such as the anterior nucleus of the thalamus *(24)*, the centromedian nucleus of the thalamus *(25)*, or the hippocampus *(26, 27)*. More specifically in the hippocampus, it has been shown that 130Hz decreases the number of seizures in patients with mesio-temporal lobe epilepsy (MTLE) *(28, 29)*. Furthermore, numerous studies in rodents have also shown the effectiveness of the 130Hz stimulation on seizures and thresholds to evoke seizures in the hippocampus *(30–32)*.

A key feature to assess the ability of a therapeutic stimulation seizures, is to analyze its impact on interictal epileptogenic biomarkers. One of the most well-known interictal biomarker of epileptogenicity is the interictal epileptiform discharges (IEDs). MTLE patients treated with HFS at 130Hz from DBS electrodes in the hippocampus had seizure frequency reduced and this was correlated with a reduction of IEDs *(33)*. A second biomarker consists in high frequency oscillations (HFO), which are divided into two groups depending on the frequency of the oscillation: 1) Ripples (approximately 150-250Hz), physiological oscillations occurring in the hippocampus during memory consolidation and learning in rodents *(34, 35)* and 2) Fast Ripples (FRs, approximately 250-500Hz) a biomarker of epilepsy *(36, 37)*. FRs are often associated with IEDs *(38, 39)*, and their surgical resection is correlated with post-surgical seizure freedom *(40)*. Similarly, DBS of the hippocampus in epilepsy can also decrease the occurrence of FRs in patients and in rodents *(41, 42)*.

Essentially, the results presented here are consistent with the above-mentioned studies demonstrating a positive effect of HFS DBS of the hippocampus. First, we performed Finite Element Method (FEM) simulations in both mice and humans. Then, by using TI from electrodes on mice’s cortex, we showed the impact on epileptic biomarkers (IEDs, FRs) with a focal 130Hz stimulation created with Pulse-width Modulated signals (PWM-TI). Along these lines, we showed the ability to scale this technique in human cadavers implanted with stereoelectroencephalography (SEEG) electrodes in order to support the possibility of using this technique as a non invasive DBS. In all applications, our PWM-TI stimulation was compared with conventional high frequency stimulation from the cortex which failed to reach deep structures and change the hippocampus epileptic biomarkers.

## RESULTS

### TI Stimulation can focally target the hippocampus, in both mice and humans

Classically, TI is obtained via the combination of two kHz sine waves *(15)* (Fig.1A). Although effective, the majority of clinical stimulation paradigms utilize square pulses for stimulation applications. In the work presented here, we aimed to demonstrate the possibility to target, in human or mice brain, and to stimulate the hippocampus with a tunable frequency. The two stimulation pairs have been oriented differently to reflect the specific orientations of the mouse hippocampus (coronal) and the human hippocampus (axial) (Fig1B). To determine where to place the cortical electrodes to evoke a stimulation in the human hippocampus, electromagnetic (EM) modeling using the finite element method (FEM) has been performed. Simulations were performed with (Supplementary Fig.1A-B) and without (Fig.1C). inclusion of SEEG electrodes (image-based placement) in the anatomical head model, for the TI stimulation and TCS. Simulations revealed that local field enhancement related to the presence of metallic SEEG contacts affect both the TCS and TI exposure - for TI exposure, an important factor is that field lines from both channels are concentrated near and perpendicularly oriented to the highly conductive material, thus maximizing TI interference. However, this effect is highly localized, affecting <1‰ of the brain volume significantly (Figure supplementary 1C; note: these simulations do not account for field pickup by the SEEG lead that could result in further field increase and power deposition near electrodes - proper lead characterization, e.g., using methodologies from *(43, 44)*, would be required to examine the stimulation and safety relevance of this effect). When comparing the TI HFS and the TCS exposure distributions, it appears that TI HFS stimulation does not impact the hippocampus overlaying cortex and brain structures as much as TCS does. Thus, we wanted to see if we could have an impact on the physiological hippocampal activities, starting with murine hippocampus.

**Figure 1:**
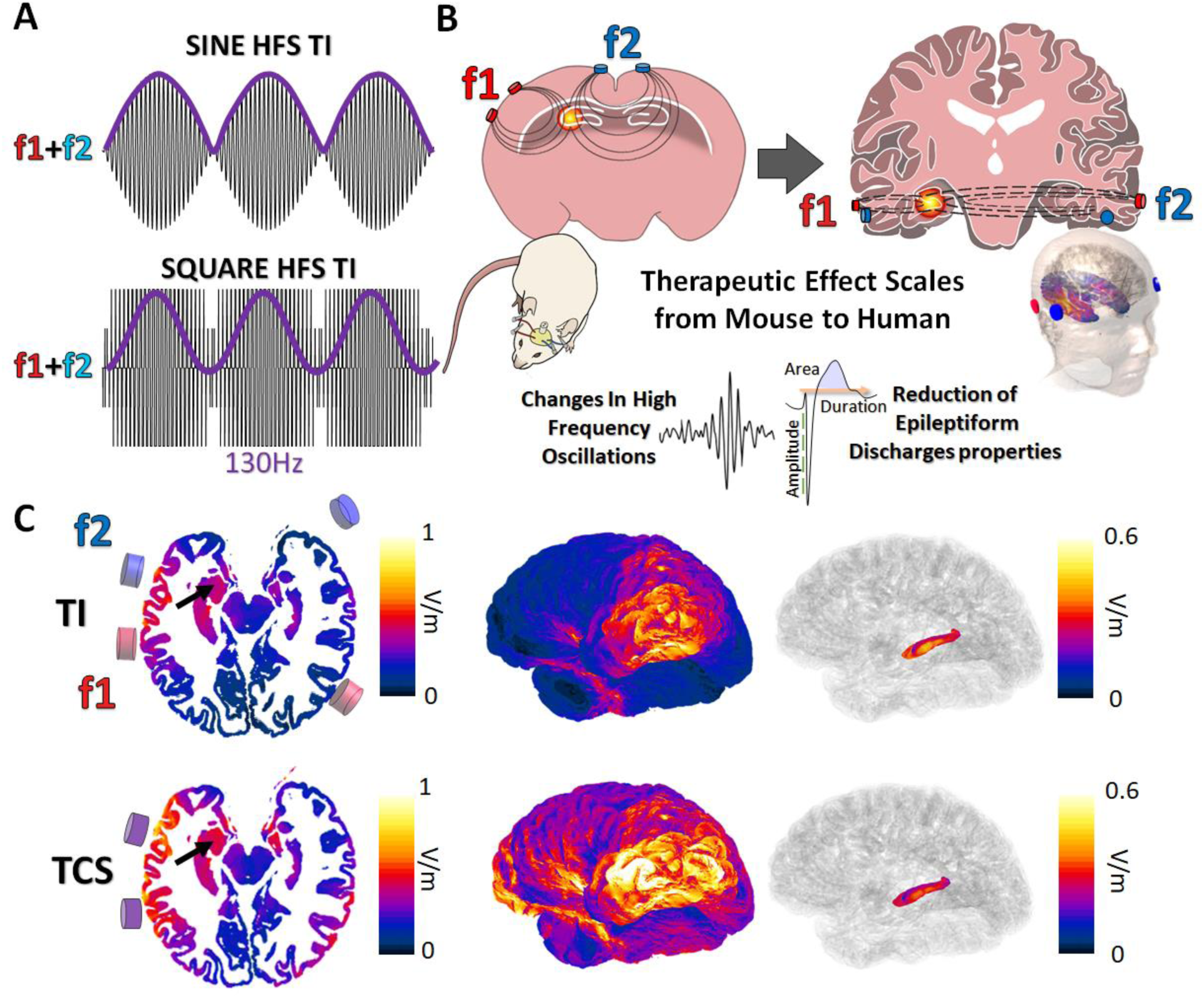
Forms of Temporal Interference and ability to scale to larger subjects. **A**. Classically, TI has been created via the combination of two kHz sine waves. Although effective, the majority of clinical stimulation paradigms utilize square pulses for stimulation applications. Thus, we investigated the impact of TI by mixing square waves (PWM-TI). **B**. Minimally invasive cortical electrodes (2 pairs) were placed on the cortex of mice and on human skin to create a minimally or non-invasive DBS via TI. In both cases, the aim was to focally reach one side of the hippocampus and to decrease epileptic biomarkers, mainly IEDs and HFOs. **C**. Simulated temporal interference envelope modulation amplitude distributions (TI; along direction of maximal modulation) and peak carrier field magnitude (bottom; TCS) and their corresponding surface field views. Arrows: hippocampus

### TI Stimulation can influence hippocampal brain rhythms in mice

In a previous study, we showed that it was possible to evoke seizures like events with PMW-TI *(45)*, the same wave was used here to perform the current study. To analyze the non-invasive DBS like effect of stimulation with TI at 130Hz (TI HFS), we compared its impact with a cortical, TCS-like, high frequency stimulation (CT HFS) and a non-treatment group (Sham) on hippocampal electrophysiological biomarkers.

To perform this, two pairs of cortical electrodes and one depth recording electrode were implanted in 29 mice. The distribution of the envelope modulation amplitude has been calculated using FEM, and compared with the field distribution of CT HFS at 130Hz. According to the simulation, the CT HFS can not activate the hippocampus, regardless which stimulation pair is selected (red shown) without activating the cortex earlier, whereas with TI HFS it is possible to stimulate the hippocampus and spare the cortex by applying two kHz square waves (f1:1300Hz f2:1430Hz) (Fig. 2A).

**Figure 2:**
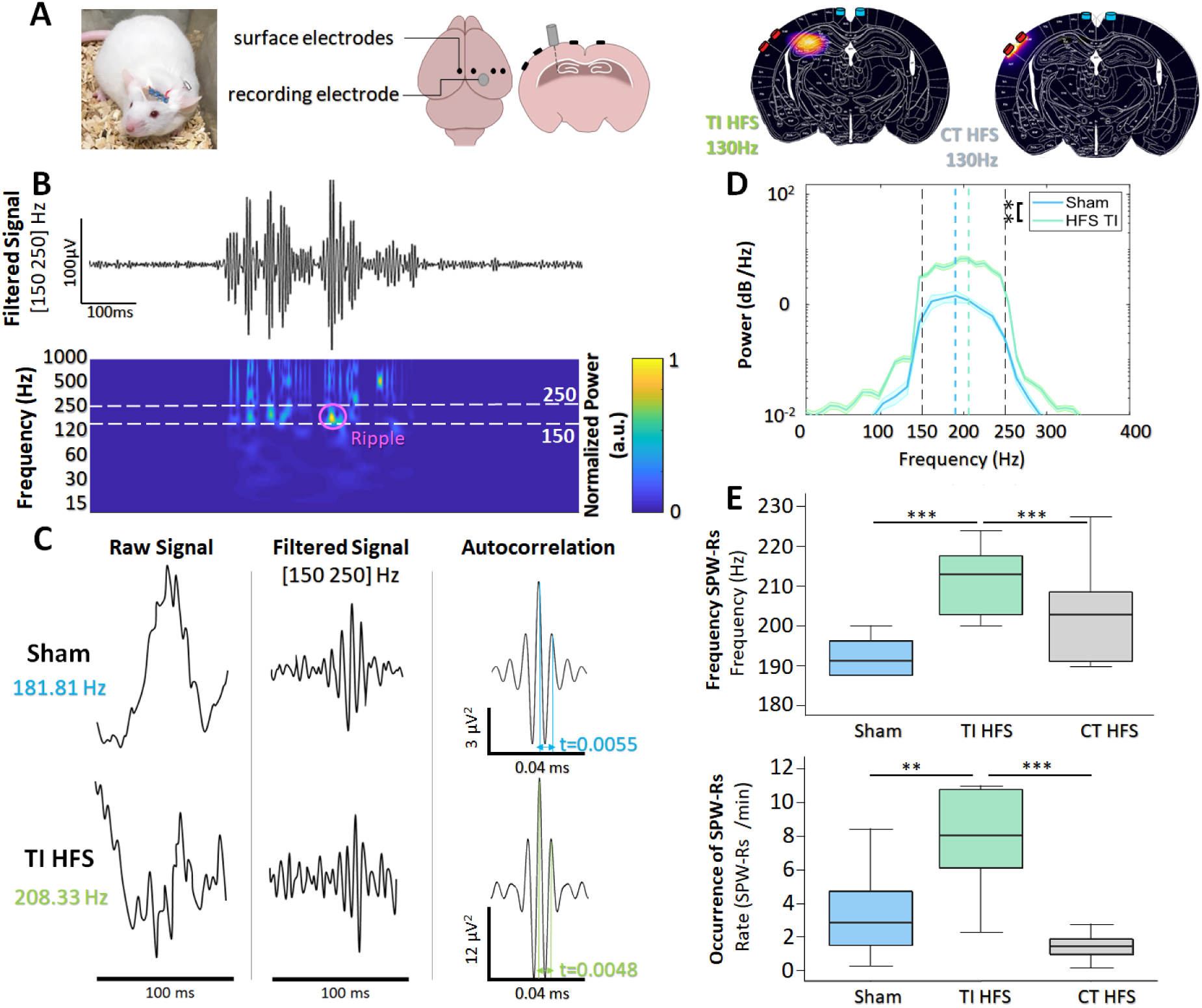
Focal hippocampal stimulation to induce SharpWave-Ripples (SPW-Rs) **A**. Mice are implanted with intracerebral electrodes to induced epileptiform activity with a kindling stimulation protocol. Finite element model of TI HFS at 130Hz and cortically applied HFS at 130Hz. It shows the creation of a hot spot of stimulation in the hippocampus via TI compared to the exclusively cortical stimulation with HFS. **B**. Raw recording and time-frequency plot of activity in the hippocampus showing the identification of SPW-Rs in the frequency band [150 250 Hz]. **C**. Example SPW-Rs and the averaged autocorrelation from Sham (spontaneous SPW-Rs, no stimulation applied) and focal TI (f1:1300Hz f2:1430Hz, envelope = 130Hz) treated mice cohorts. The distance in time of the first and second peak of the average autocorrelation shows a clear increase in SPW-Rs’ frequency. **D**. The power spectral density of SPW-Rs also shows the shift in SPW-Rs’ frequency and additionally the increase in SPW-Rs’ strength. **E**. The shift in frequency is not seen for HFS treated mice, where 130 Hz were applied directly to the cortex. Focal TI stimulation causes a direct increase in the incidence of SPW-Rs (SPW-Rs/min).

Hippocampal sharp wave–ripple complexes (SPW-Rs) are observed in both slow-wave sleep and in awake behavior during inactivity and are considered to be involved in the transfer of stored information to the neocortex for memory consolidation *(46)*. SPW-Rs are oscillations that can be detected in time frequency plot, where they are creating a frequency region of activation in the [150 250] Hz band (Fig. 2B). Behrens et al. first demonstrated that it is possible to evoke SPW-Rs via an implanted device and that their properties change depending whether SPW-Rs are physiological or induced *(47)*.

Here, we observed that there are also differences between the properties of intrinsic SPW-Rs (sham) and focally induced (TI HFS) SPW-Rs (Fig. 2C). The averaged autocorrelation of SPW-Rs from the two cohorts of mice allowed us to determinate the frequency of the detected SPW-Rs. We demonstrated by calculating the distance in time between the first and second peak of the autocorrelation (Sham: t=0.0055s TI HFS: t=0.0048s) a clear shift in the SPW-Rs’ frequency induced with focal TI HFS (from respectively 181Hz to 208Hz). In addition, the power spectral density of the average (more than 1000 detected SPW-Rs and induced SPW-Rs) showed also a shift in SPW-Rs’ frequency. However, it was also observed a significant increase in SPW-Rs strength for induced SPW-Rs, again as expected from a focal stimulation of the hippocampus (Fig. 2D) (*Sham vs TI HFS p-value=0*.*003)*. The CT HFS stimulation had clearly not the same impact on SPW-Rs, where SPW-Rs’ frequencies and occurrence for Sham and CT HFS were equivalent (*Frequencies: Sham vs TI HFS: p-value < 0*.*001 – Sham vs CT HFS: p-value = 0*.*073, CT HFS vs TI HFS: p-value < 0*.*001)*. This may indicate that the SPW-Rs analyzed were physiologic and not induced by the stimulation. Additionally, only TI HFS created the expected increase in SPW-Rs’ incidence (SPW-Rs/min), where TI HFS can multiply by 4 the number of SPW-Rs detected (Fig. 2F) (*Occurrence: Sham vs TI HFS: p-value = 0*.*003, CT HFS vs Sham: p-value = 0*.*278, TI HFS vs CT HFS: p-value < 0*.*001)*.

Taken together, these results indicated the ability of TI HFS at 130Hz to achieve a highly focal stimulation of the hippocampus, equivalent to previous studies evoking similar results with direct “in situ” depth stimulation. Such effect was not observed with a transcranial stimulation without TI (CT HFS). We then analyzed the anti-epileptic effect of our TI HFS by recording the number of IEDs and the occurrence of FRs [250 500 Hz] in a mouse model of epilepsy.

### TI therapy: HFS TI at 130Hz decreases the expression of epileptogenic biomarkers

As mentioned above, IEDs and FRs are well-known biomarkers of epilepsy in both humans and animal models *(48)*. To assess the influence of TI stimulation, four cohorts were considered: Baseline (mice before any treatment), Sham (mice electrically kindled to have epilepsy), TI HFS (mice electrically kindled receiving HFS focal TI at 130 Hz in the hippocampus), and CT HFS (mice electrically kindled receiving direct 130 Hz from the cortex surface). Baseline recordings were made in all mice before undergoing the kindling protocol (Fig. Supplementary 2). The kindling lead mice to express epileptogenic biomarkers, like pathological IEDs *(49)*. Traditional analysis of IEDs evaluates several components such as rate, area under the wave, duration of the IEDs, and amplitude (Fig. 3A) *(50)*. For rate, IEDs recorded after the TI HFS 130Hz treatment occurred less frequently (decreased by a factor of 3 compared with Sham). Only TI HFS was able to reduce the number of IEDs to a level similar to before the induction of epilepsy (Baseline) *(<u>significant:</u> sham vs TI 130Hz: p-value = 0*.*001; Baseline vs CT HFS 130Hz: p-value = 0*.*041; Baseline vs Sham* : *p-value = 0*.*005 / <u>non-significant :</u> CT HFS vs TI 130Hz* : *p-value = 0*.*384; CT HFS 130Hz vs sham* : *p-value = 0*.*401; Baseline vs TI 130Hz: p-value = 0*.*287)* (Fig. 3B).

**Figure 3:**
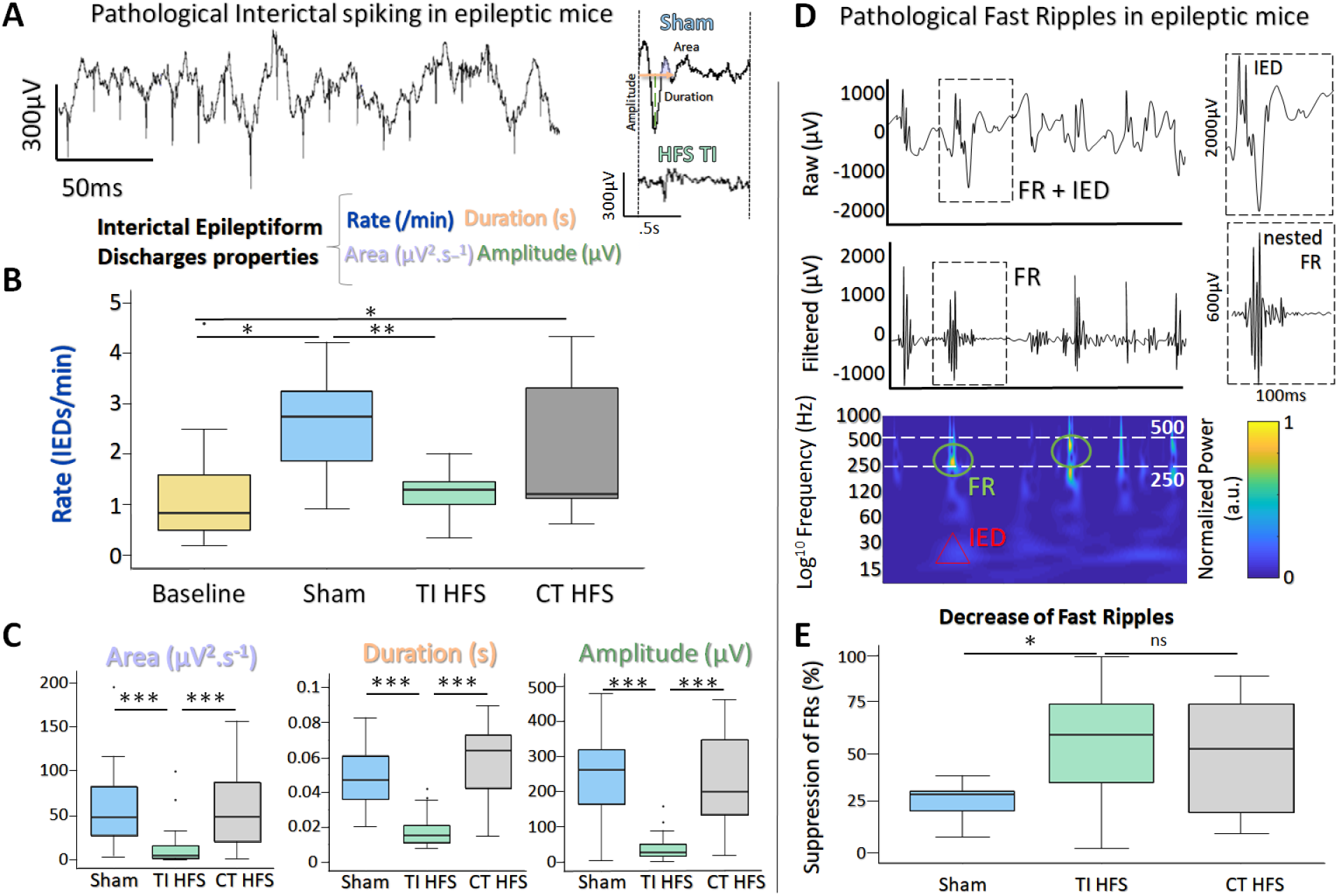
Suppression of Interictal Epileptiform Discharges (IEDs) and Fast Ripples (FRs) with PWM-TI. **A**. Pathological IEDs and their properties: Rate, Duration Amplitude and Area of the wave. **B**. Analysis of the IEDs rate before and after the treatment. Focal TI HFS of the hippocampus dramatically reduces pathological IEDs levels and can reproduce the baseline (before kindling) stage; however, traditional stimulation of the cortex (HFS 130Hz) has little effect, and cannot decrease the IEDs rate to pre-kindling levels. **C**. Analysis of the IEDs properties. Area, Duration and Amplitude were calculated for the detected IEDs. Only TI HFS can reduce the area of the wave (5 times smaller), duration of the IEDs (3 times shorter) and the amplitude (2 times lower). **D**. Raw and filtered [250 500] Hz signal with detected FRs. Time frequency plot showing the activation in the FRs band. **E**. Pathological FRs are seen to also decrease with focal TI HFS, with a 50% suppression of the FRs after the treatment. A ratio for each mice was calculated to compare pre- and after-treatment. There is a 25% diminution of FRs when no treatment was applied, which shows the impact of the time on these biomarkers. TI HFS is the only treatment allowing a significant decrease of FRs compared to Sham.

For the analysis of the characteristics of IEDs, TI HFS treatment significantly decreased the area under the wave, the total duration, and the amplitude of the IEDs (<u>Area</u> - *Sham vs TI 130Hz: p-value < 0*.*001; CT HFS 130Hz vs TI 130Hz* : *p-value < 0*.*001; CT HFS 130Hz vs sham* : *p-value = 0*.*761 -* <u>Duration</u> - *sham vs TI 130Hz: p-value < 0*.*001; CT HFS 130Hz vs TI 130Hz: p-value < 0*.*001* ; *HFS 130Hz vs sham* : *p-value = 0*.*183* - <u>Amplitude</u> - *sham vs TI 130Hz: p-value < 0*.*001; CT HFS 130Hz vs TI HFS 130Hz* : *p-value < 0*.*001; CT HFS 130Hz vs sham* : *p-value < 0*.*001*).

In all cases, TI HFS treatment of mice gave rise to a significant reduction of epileptic biomarkers, while CT HFS and Sham were unable to achieve such effects (Fig. 3C).

The second epileptogenic biomarker analyzed was FRs, an indicator of epileptogenicity used in the identification of the epileptogenic zone *(51)*. FRs are oscillations which can be detected after filtering the raw signal, typically between [250 500 Hz]. Interestingly, FRs very often co-occur with IEDs which has been stated as a good indicator for epileptogenicity *(52)* (Fig. 3D). We analyzed the change in FRs, a ratio before and after treatment (or no treatment in the case of sham) was performed. For TI HFS, a clear decrease in the number of FRs was seen with more than 50% FRs suppression. For CT HFS stimulation, there was no significant impact on the reduction of FRs compared to Sham even if there was no significant differences compared to TI HFS (Fig. 3E). (*Sham vs TI 130Hz: p-value= 0*.*023; CT HFS 130Hz vs TI 130Hz: p-value= 0*.*327; CT HFS 130Hz vs sham: p-value = 0*.*132*).

Overall, these results illustrated the impact of the TI HFS at 130Hz, upon the two biomarkers of epilepsy within the hippocampus, reducing both the features and the number of IEDs, and the prevalence of FRs.

### Scaling HFS TI: from the mouse to the human brain

Here, we explored the possibility of applying non-invasive TI stimulation targeting the hippocampus in human by analyzing the HFS TI stimulation in human cadavers. Several SEEG electrodes were implanted in the temporal lobe of cadavers to record the applied stimulation potential (Fig. 4A). By mixing 1300Hz and 1430Hz with electrodes placed on the skin, a focus of stimulation in the anterior part of the hippocampus was created as previously done in mice. The amplitude of the envelope of stimulation was at 8mV only in the hippocampus. The two other electrodes placed in the central area (electrodes c &f) recorded also an envelope amplitude around 5mV. Electrodes placed outside of the hippocampus and contralateral SEEG electrodes didn’t record any amplitude modulated signal (Fig. 4B). This result confirmed that TI can reach deep-seated structures, here the hippocampus, in human cadavers, and shows that scaling of the TI HFS as previously done in mice is achievable in a human subject. To further analyze the effect of the PWM signal used across the study (TI HFS), we compared its effect with the traditional TI with sine waves. We show an example electrode (f) to illustrate the main differences between the two forms of TI and additionally include transcranial stimulation (TCS) at 130Hz using sine waves. The modulation envelope amplitudes increase at depth in the cases of square and sine TI as is shown on the raw recording in the bottom right panels (Fig 4 C). Between the contacts n°8 (superficial) and n°2 (deepest), the amplitude of the envelope (upper part of the recorded signal) is multiplied by 4. Looking at the TCS 130Hz, only the cortex gets stimulated, and the amplitude of the signal decreases with depth.

**Figure 4:**
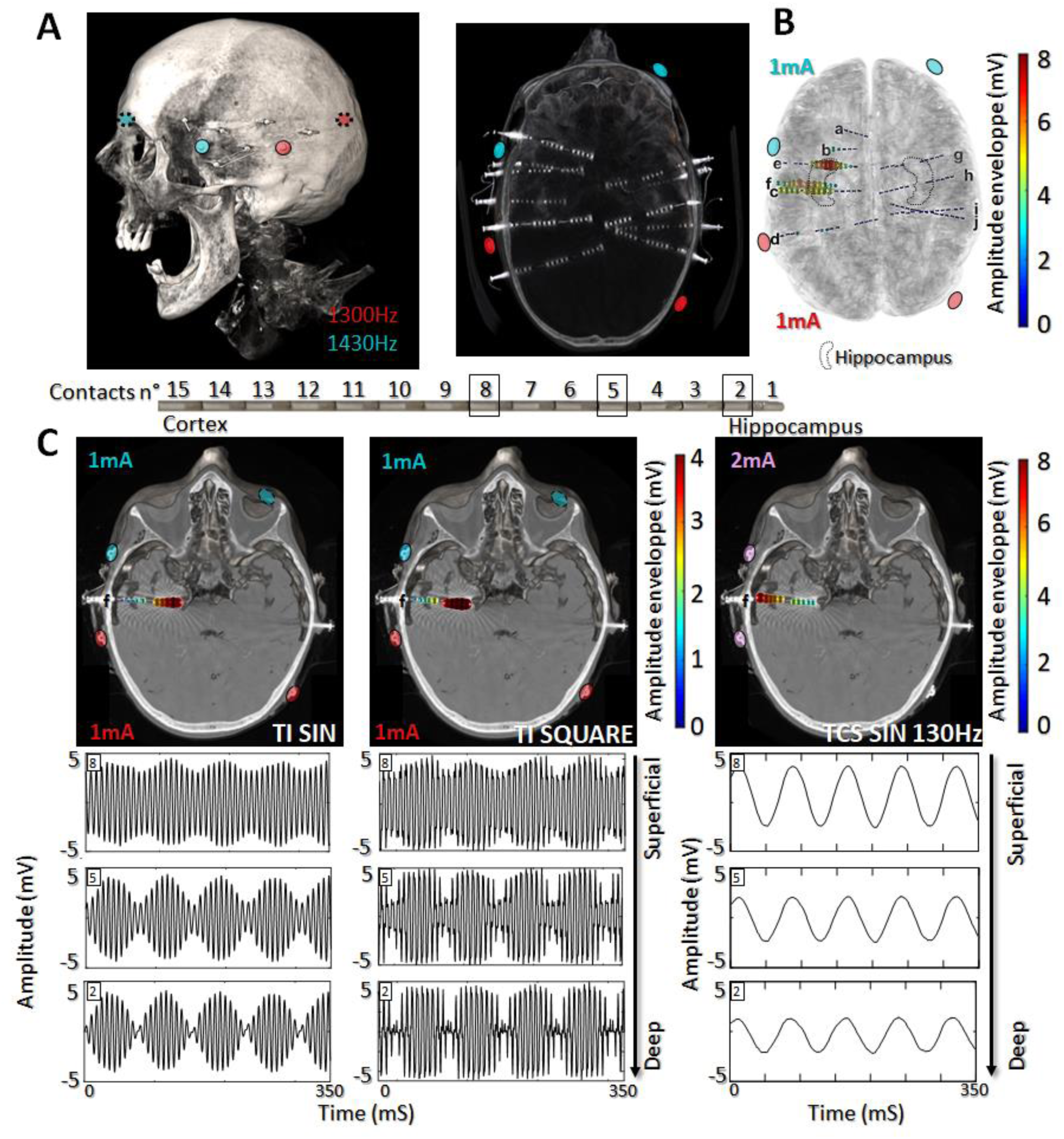
Scaling the effect of the HFS TI at 130Hz in the human head: **A**. Placement of the electrodes on the skin of the human cadaver. Skin electrodes were used to deliver the TI HFS 130Hz in both square and sine wave. 10 Stereotaxic electro-encephalographic electrodes (SEEG) were implanted in the brain as for a MTLE patients in order to record the stimulation potentials inside the brain. **B**. Co registration with the scanner and a template MRI show the amplitude of the envelope recorded during TI HFS 130Hz session. It gives an indication on where the focus of stimulation is and the amplitude of the envelope of stimulation. Another advantage of the TI HFS is to move the maximum of the envelope without moving the skin electrodes. Here, 1mA and 3mA were chose to better target the anterior hippocampus in this specific cadaver and the field modulation envelope can reach an amplitude of 7mV. **C**. the SEEG electrode f is used to show the depth of the focal stimulation of TI HFS in both square and sine waves. For transcranial (TC) HFS 130Hz, it shows stronger stimulating fields in the cortex, which is primarily activated, compared to the deep structures, unlike for TI with sine waves or square waves (PWM TI).

In our human cadaver experiments, we used SEEG electrodes to record the potential created by the HFS TI at 130Hz. We showed that by stimulating with TI, a significant envelope amplitude is recorded at depth, and it decreased on the electrodes’ contacts placed in the cortex. This is not the case when 130Hz is applied via TCS where the largest amplitude is recorded at shallow.

## DISCUSSION

For patients with intractable epilepsy not eligible for surgery or with failure of previous resective surgery, neuromodulation techniques turn out to be promising options *(24, 53)*. These techniques encompass invasive (*e*.*g*. DBS) and non-invasive stimulations (*e*.*g*. TCS). Invasive techniques are costly, require invasive surgery (implantation of permanent depth electrodes) and are at risk of complications. A complementary non-invasive stimulation which could provide similar results would be undoubtedly very useful. Some TCS methods are already promising and show some degree of seizure reduction, particularly those using tDCS *(54)*. However, TCS is probably limited by the depth of penetration and spread of the applied electric fields The results of this study appear encouraging and prompt to develop and refine TI as a new TCS paradigm capable of modulating the activity of deep structures (hippocampus in this study).

Mouse brain stimulation experiments in this work have demonstrated that superficial cortical TI HFS, but not CT HFS, is able to modulate physiological and pathophysiological activities of the hippocampus. For healthy physiological activity, it is understood that focal electrical stimuli can induce long-term potentiation (LTP), the neurophysiological process underpinning learning and memory, and will lead to the generation of SPW-Rs in the hippocampus *(55)*. In mice, TI HFS can evoke SPW-Rs, which have higher oscillation frequencies compared to the intrinsic SPW-Rs recorded in Sham conditions, replicating the results of focal stimulation of the same hippocampal region with depth-electrodes. In contrast, CT HFS applied on the cortex was not able to modify the hippocampus activity. Here, we showed an ability of the focal TI to influence natural brain rhythms in the form of SPW-Rs and induced SPW-Rs, shifting central frequencies, strength, and incidence of SPW-Rs, never seen with non-invasive direct hippocampal stimulation.

For pathophysiological activity in mice, TI HFS was shown to decrease both FRs and IEDs. In the case of IEDs, the spike rate returned to baseline (before epilepsy induction) and can also significantly reduce epileptic features of IEDs (amplitude, duration, etc.). Traditional transcranial methods of stimulation are not completely ineffective, and in the study here we see that HFS applied to the cortex has an effect sized between TI HFS and Sham. CT HFS is not as focal and its effect is not as dramatic, which is line with the literature *(13)*.

The results in mice have then been scaled to humans. In our human cadaver experiments, we used SEEG electrodes to record the potential created by the TI HFS at 130Hz. We clearly demonstrated that we obtained larger electrical amplitudes in depth when using TI and reduced stimulation amplitudes in the surrounding cortex. This is not the case when 130Hz is applied via conventional TCS stimulation. We showed that the cortex near the electrodes has the largest amplitude of stimulation, and that the recorded electric field unsurprisingly decreases with depth. However, the main difference between our rodent and human experiments are the invasiveness of the stimulating electrodes. In mouse experiments, the position of the electrode was minimally invasive: pairs of electrodes were located on the cortex surface to avoid any motion during freely-moving-awake behavior. For humans, skin electrodes were used on the scalp, creating a fully non-invasive deep brain stimulation. For long term implants, we imagine a subcutaneous implant to increase the applicable current by avoiding possible tingling sensations, to reduce current shunting through the scalp, and to be more adapted for long-term stimulation. It would be also necessary that the stimulation electrodes be firmly screwed to the bone at the level of the outer table and be maintained perfectly immobile.

In summary, TI, as a completely non-invasive stimulation method, was able to achieve deep brain structure (hippocampus). As such it offers new perspective of non-invasive neuromodulation techniques for patients with refractory epilepsy ineligible to surgery or with failure of the latter and need of stimulation of deepest brain structures. TI could significantly change the manner in which neuromodulation in epilepsy is delivered, because it can be no longer necessary to receive an invasive implant. A minimally invasive form of TI therapy could be prescribed (*e*.*g*. subcutaneous electrodes) or alternatively an intermittent, totally non-invasive therapy could be tested for efficacy before a potential DBS device implantation.

## MATERIALS AND METHODS

### EM exposure modeling

Electric exposure simulations have been performed using the structured ohmic-current-dominated electro-quasistatic finite-element-method solver from Sim4Life (ZMT Zurich MedTech AG, Switzerland), which has been verified to suitably approximate Maxwell’s equations for the frequencies and setup of interest (Fig.1). The highly detailed MIDA anatomical head model *(56)* that distinguishes >130 different anatomical regions was used, and the two applied currents were simulated individually by applying corresponding Dirichlet voltage boundary conditions to the respective electrode pairs and normalizing to the resulting total current. Electrical conductivity values were assigned to the different tissues *(57, 58)*. To simulate the impact of the SEEG presence in terms of local field enhancement due to the presence of highly conductive metal contacts, CT images from a patient were co-registered with the MIDA model using Sim4Life and ensuring a good match between skulls. Thresholding of the metal-related image-artefacts in the CT scan was used to guide the placement of 14 SEEG lead models featuring 13-17 cylindrical electrode contacts each (modeled as perfect electric conductors) separated by insulating segments according to manufacturer specifications. Rectilinear discretization with a resolution of 0.27-0.65mm (maximal refinement at the SEEG contacts) was employed (142 million voxels).

The following quantities were extracted for analysis and visualization using native Sim4Life postprocessing functionalities, as well as custom Python scripts: maximal TI modulation amplitude and peak high-frequency (HF) field magnitude according to the equations from *(15)* their normal component on the brain surface (as a measure for cortical TI and HF stimulation), overall and hippocampal peak, peak 2mm-averaged (according to ICNIRP Guidelines on the exposure safety standard *(59)*), and isopercentile values.

### Animals

All experiments were performed in accordance with European Council Directive EU2010/63, and French Ethics approval (Williamson, n. APAFIS#20359-2019041816357133 v10). Animals were kept in transparent cages in groups of three to five, in a temperature-controlled room (20 ± 3°C) with a 12/12h night-day cycle. All animals had *ad libitum* access to food and water.

### Surgical Procedure

For this study we used 29 male OF1 mice (Charles Rivers Laboratories, France) aged 8-10 weeks. Animals were kept in transparent cages in groups of three to five, in a temperature-controlled room (20 ± 3°C) with a 12/12h night-day cycle. Mice were implanted with two pairs of minimally invasive cortical electrodes with an intracerebral depth-electrode in the hippocampus. The hippocampal depth-electrode provided the induced epileptic activity *via* a standard kindling protocol, and subsequently remained in place to record the electrical activity of the hippocampus. Mice were divided into groups, 12 mice received the TI HFS treatment, 9 others were sham (they underwent the surgical procedure and the kindling protocol but no treatment) and 8 mice had the 130 Hz cortical stimulation (CT HFS).

Mice were anaesthetized *via* an intraperitoneal injection of ketamine (50mg/kg) and xylazine (20mg/kg) and placed in a stereotaxic frame with the head adjusted for bregma and lambda in the same horizontal plane. After midline scalp incisions, the following stereotaxic coordinates were used for craniotomies: Cortical electrodes [AP: -1.94, ML: +0.5; -0.5; -3.9; -4.3] Implantable twisted-pair platinum electrodes (from PlasticsOne; wire length = 5mm, individual wire diameter = 125µm) [AP: -2.7, ML: +2.04, DV: 1.30] using a 20-degree angle to reach the hippocampus by considering the constraints due to the location of the minimally invasive cranial electrodes. All the coordinates were calculated using the Paxinos Atlas. The four stainless steel mini-screws (Component Supply, Miniature Stainless Steel Self-Tapping Screws: TX00-2FH) were placed on the cortex without penetration into the brain tissue. Subsequently, dental cement (Phymep, SuperBond) was applied on the skull surface to fix the screws and the skull cap was formed using Dentalon. After the surgery, all mice were kept in separate cages to avoid fighting and to avoid damage to implanted electrodes. During the post-surgical recovery time (7 days), all the mice were observed for signs of pain, distress and neurological complications.

### Kindling protocol

After 7 days of recovery following surgery, a protocol to induce and to characterize the afterdischarge threshold was applied using the Racine scale by the implantable electrode. Following previous work, ADs were defined as high-amplitude IEDs and polyspike epileptiform events visible after the applied stimulus. We used a 10 second stimulation, bipolar, biphasic, at 50 Hz with a 500µsec pulse width, with the amplitude increased in 50 µA steps, starting from 50 µA, until reaching the seizure of stage 4 on Racine scale. After the determination of the AD threshold, the parameters were used to do the kindling protocol where 10 stimulations were performed with 10 minutes of rest between each stimulation. 24 hours after, the same protocol was repeat to evoke spontaneous epileptiform events. Two baselines were recorded, before and after the kindling protocol.

### Electrical stimulation

*In vivo experiments:* At day 8 after surgery, 1 hour pre-therapy recordings were made, followed by a 1-hour therapy using either Sham (No stimulation), HFS TI (TI with 130Hz envelope) or CT HFS 130Hz (standard transcranial stimulation of 130Hz applied to the cortex), followed by a 1-hour post-therapy recording. The two frequencies applied to create the 130Hz stimulation is f1: 1300Hz & f2: 1430Hz. Square pulses were used to perform the stimulation, like used in clinic, called here PWM-TI. Treatment was applied with a function generator (keysight technology, serie edu33212a) at 1300Hz and 1430Hz (the two sources are independent). For the TI treatment, the two frequencies created a focal 130 Hz envelope frequency in the hippocampus. The square TI used biphasic pulses of 100 μs with pulse amplitudes of 60%*AD threshold μA for one hour. For the HFS treatment, 130 Hz has been directly applied by one pair of the cortical electrodes with the same amplitude and duration as TI HFS treatment. For the sham condition, no treatment has been applied, however animals were recorded for the same amount of time as TI and cortical treatment.

### Recordings

*In vivo experiments:* All recordings have been performed by electrical stimulators (IntanTech, Intan 128ch Stimulation/Recording Controller) with a sampling rate at 30KHz. To process the data, all files have been converted from RHS format to a readable format for us using Matlab format and specific format of our homemade software AnyWave *(60)*.

### Human cadaver

Anatomical subjects were provided by the “*service des corps donnés à la science*” by Aix Marseille University and all the experiments were performed in the Faculty of Medecine *La Timone* (Aix Marseille University). All the subjects were perfused with zinc chloride and stored in freezer until the experiments. Subjects were left at least 2 hours at 20°c before any stimulation / recording session. For depth recording, SEEG electrodes (Alcis©, France) and for scalp stimulation, classical ECG electrodes were used (Ambu®). All the SEEG insertions were performed by a neurosurgeon (RC) on the basis of his surgical expertise of SEEG for adult patients (robotized and frame-based procedures) but the implantation was based on mere anatomical landmarks on entry point without stereotactic reference. For electrode localization, a CT scan of the head was performed after the stimulation experiments at CERIMED, Aix Marseille University, France. All recordings were done with RHS recording and stimulation controllers (Intan®) and stimulation was done by either the RHS recording and stimulation controller or a function generator (Keysigth ®) driving a DS5 current source (Digitimer ©) to reach the desired current.

### Analysis

#### In vivo experiments

*To* detect specific events as IEDs and High Frequency Oscillation (HFO), we performed a semi-automatic detection on the signal. The automatic detection part was processed by using Delphos software, a detector of IEDs and oscillations used mainly in clinical research. *(52)* Delphos is based on a method of whitening (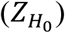) the time-frequency spectrum to optimize the signal to noise ratio at each frequency. Events of interest are detected while they are above a specific threshold in the spectrum. Before to apply Delphos on the signal, it was down sampled at 3750Hz. As Delphos has mainly been apply on clinical data, parameters were changed to fit with the mice data. Once events detected, they were reviewed by an expert (EA) by using AnyWave software, a visualization software for electrophysiological data. *(60)* The events were accepted/rejected by the expert. Finally, the events (IEDs [15 45]Hz, SPW-Rs[150 250]Hz and FRs[250 500]Hz) were extracted in Matlab format to perform an average of all events and determine the duration, the amplitude and the rates by minutes.

#### Human cadaver

To ensure the correct anatomical position of the SEEG electrodes (possibility of deviation in their trajectories), especially in the hippocampus, a post-operative CT-scan of the cadaver’s head was performed. Then we used Gardel, to localize and label SEEG contacts. *(62)* Gardel requires an MRI image (T1w) and CT scan done after the implantation. For the T1w image, we used the MNI template as we didn’t have one for the cadavers. MRI image was co-registered on CT image to work on CT space. Then, the electrodes were automatically detected and reviewed and labeled manually. This method enabled us to extrapolate as to where the electrodes would be in a living subject with a normal non-shrunk and atrophic brain. To analyze the intra-cerebral EEG signal, as for mice, the data was first converted from RHS to Matlab format. We used Matlab, especially the Signal Processing toolbox, to determine the envelope of stimulation induced by the TI at 2000Hz and 2005Hz. Firstly, the signal was filtered with band pass filter with a passband frequency of [1000 3000] for TI. Secondly, the magnitude of its analytics signal (envelope) was calculated on the filtered signal by using Hilbert transformation on the filtered signal. A sliding window of 230ms was used to determine the amplitude of the envelope peak-to-peak in mV, by subtracted the minimum of the envelope to the maximum. The median value of all the window was used as amplitude value for the contact. This process has been performed for each contact which allows to have an amplitude value by contact. Finally, the amplitude value was projected on the electrodes in the mesh of the anatomical image (MNI template) to visually determine which part of the brain received the stimulation.

### Statistical analysis

#### In vivo recordings

This study has been designed so that all statistical tests have a power of 80%. After gathering all LFP recordings for all mice (n=29), FRs, IEDs and SPW-Rs were analyzed using R studio®. In order to adapt the test to the data distribution, all groups were tested for normal distribution using Shapiro tests. As distributions were not normal, Wilcoxon rank tests were performed to highlight differences in between the different groups (Alpha = 5%). Regarding physiological ripples, occurrence and in our three conditions (Sham, TI HFS and HFS) were compared. For pathological fast ripples, the suppression of these epileptic markers has been calculated using a ratio (PostTreatment*100/PreTreatment) to show the effect of our treatment groups. Finally, IEDs were analyzed in the same manner whether they were recorded in mice or human: Rate, Duration, Area of the wave and Amplitude of the IEDs were calculated and compared among our three groups.

## Acknowledgments

We would like to thank Mme Poulin E. and Dr Serratrice N. for their help regarding experiments with human cadavers in Service des corps donnés à la science.

## Funding

This research was supported by funds from the European Research Council (ERC) under the European Union’s Horizon 2020 research and innovation programme (grant agreement No. 716867).

## Author contributions

A.W. conceived the project with input from N.G. and E.A.. E.A., S.L., C.L., P.D., F.M. and B.B. performed experiments. B.B. E.N. and A.C. performed finite-element simulations. E.A. and A.J. analyzed neural data. A.W. and E.A. wrote the paper with input from the other authors, including R.C., F.B. and V.J.

## Competing interests

The authors declare no competing financial interests.

## Supplementary Figures

**Supplementary figure 1:**
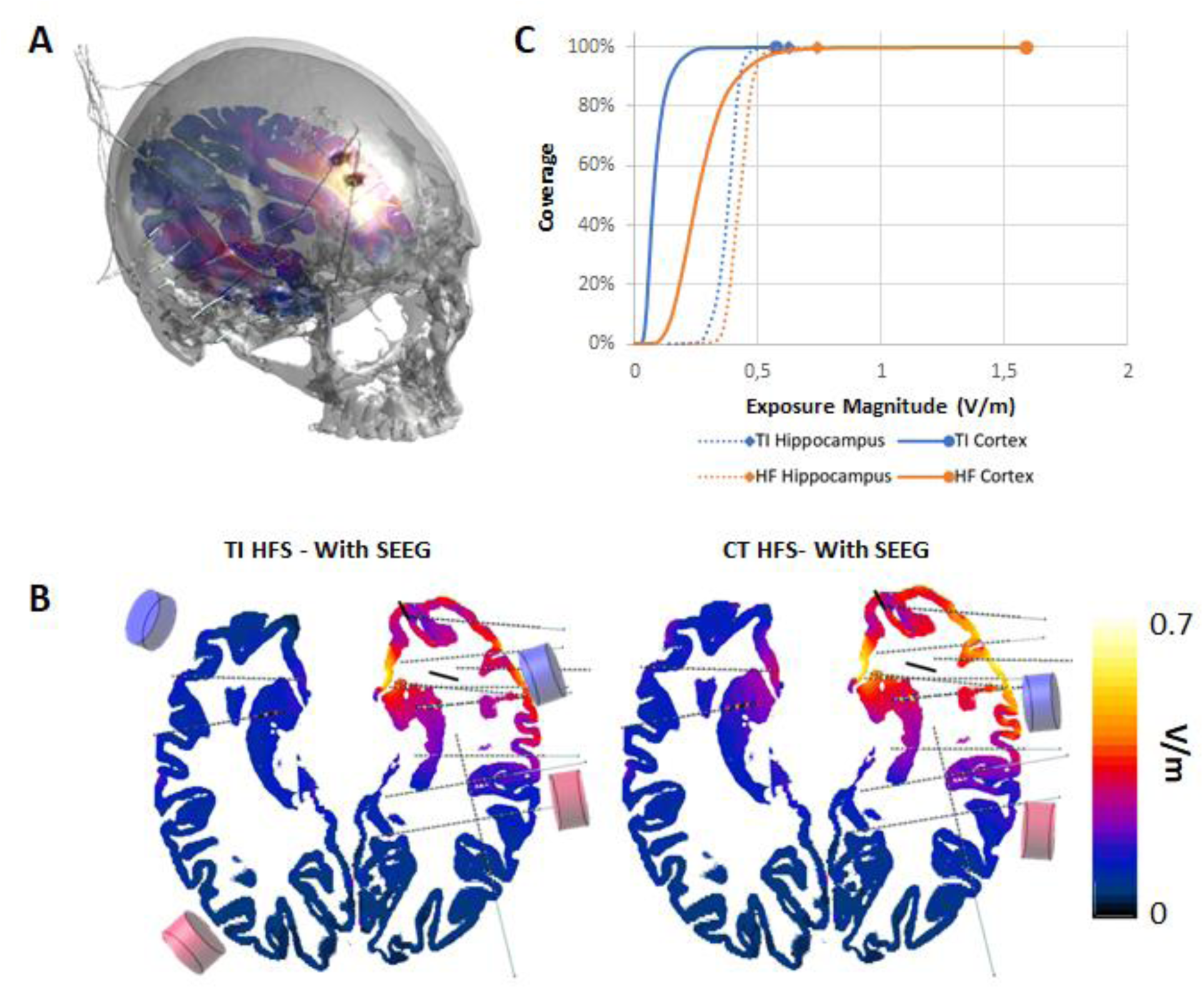
EM simulations reveal highly localized field enhancement by the presence of metallic SEEG electrodes. **A**. illustration of the head models with the SEEG leads and the emplacement of the stimulation electrodes **B**. Simulated temporal interference envelope modulation amplitude distributions (left: TI along direction of maximal modulation) and peak carrier field magnitude (right) in the presence of the SEEG leads, at a normalized current of 1mA per channel. **C**. cumulative histograms of the cortical and hippocampal TI modulation and peak carrier distributions.

**Supplementary figure 2:**
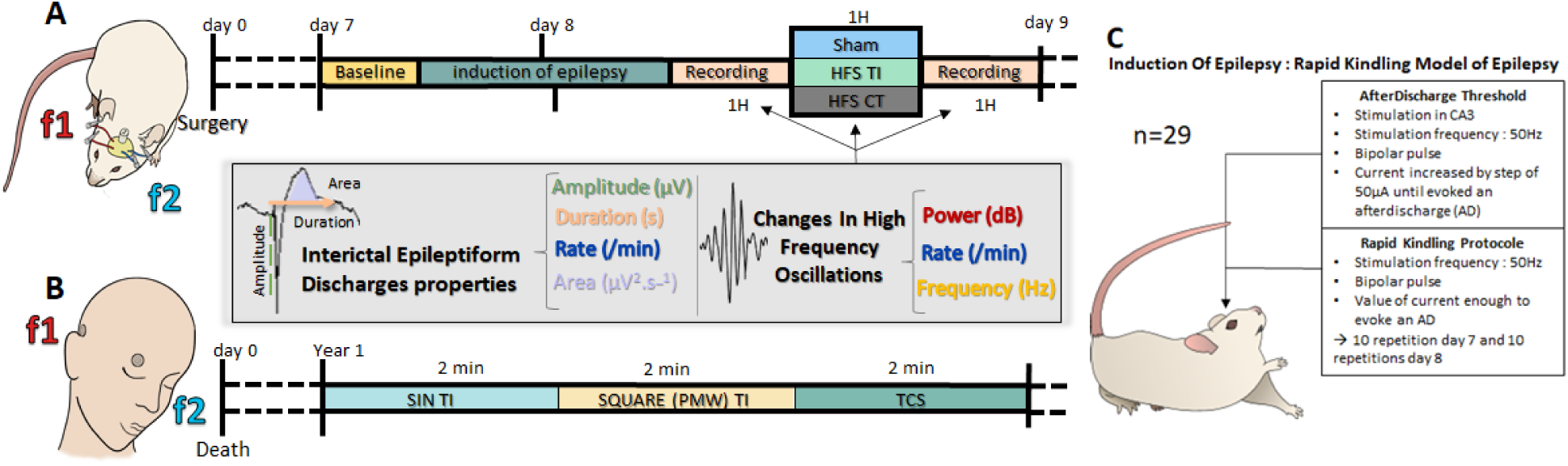
Application of TI on human and mice. **A**. Epilepsy in mice has been induced via a kindling model. Then PMW TI has been applied for one hour and analysis has been performed on the epoch before and after the TI HFS. **B**. In cadavers, TI (sin and square at 130Hz) has been applied for 2 minutes each. **C**. Characteristics of the rapid kindling model of induction of epilepsy used.

## Notes

### Competing Interest Statement

The authors have declared no competing interest.

## References and Notes

1. K. M. Fiest, K. M. Sauro, S. Wiebe, S. B. Patten, C.-S. Kwon, J. Dykeman, T. Pringsheim, D. L. Lorenzetti, N. Jetté, Prevalence and incidence of epilepsy: A systematic review and meta-analysis of international studies. Neurology 88, 296–303 (2017).

2. W. Löscher, D. Schmidt, Modern antiepileptic drug development has failed to deliver: ways out of the current dilemma. Epilepsia 52, 657–678 (2011).

3. P. Ryvlin, J. H. Cross, S. Rheims, Epilepsy surgery in children and adults. Lancet Neurol. 13, 1114–1126 (2014).

4. A. L. Velasco, F. Velasco, M. Velasco, D. Trejo, G. Castro, J. D. Carrillo-Ruiz, Electrical Stimulation of the Hippocampal Epileptic Foci for Seizure Control: A Double-Blind, Long-Term Follow-Up Study. Epilepsia 48, 1895–1903 (2007).

5. R. S. McLachlan, S. Pigott, J. F. Tellez-Zenteno, S. Wiebe, A. Parrent, Bilateral hippocampal stimulation for intractable temporal lobe epilepsy: Impact on seizures and memory. Epilepsia 51, 304–307 (2010).

6. A. E. L. Warren, L. J. Dalic, W. Thevathasan, A. Roten, K. J. Bulluss, J. Archer, Targeting the centromedian thalamic nucleus for deep brain stimulation. J. Neurol. Neurosurg. Psychiatry 91, 339–349 (2020).

7. K. Ashkan, P. Rogers, H. Bergman, I. Ughratdar, Insights into the mechanisms of deep brain stimulation. Nat. Rev. Neurol. 13, 548–554 (2017).

8. N. Zangiabadi, L. D. Ladino, F. Sina, J. P. Orozco-Hernández, A. Carter, J. F. Téllez-Zenteno, Deep Brain Stimulation and Drug-Resistant Epilepsy: A Review of the Literature. Front. Neurol. 10, 601 (2019).

9. M. L. Kringelbach, N. Jenkinson, S. L. F. Owen, T. Z. Aziz, Translational principles of deep brain stimulation. Nat. Rev. Neurosci. 8, 623–635 (2007).

10. M. C. H. Li, M. J. Cook, Deep brain stimulation for drug-resistant epilepsy. Epilepsia 59, 273–290 (2018).

11. J. G. Pilitsis, Y. Chu, J. Kordower, D. C. Bergen, E. J. Cochran, R. A. E. Bakay, POSTMORTEM STUDY OF DEEP BRAIN STIMULATION OF THE ANTERIOR THALAMUS: CASE REPORT. Neurosurgery 62, E530–E532 (2008).

12. P. Sudbrack-Oliveira, M. Z. Barbosa, S. Thome-Souza, L. B. Razza, J. Gallucci-Neto, L. da Costa Lane Valiengo, A. R. Brunoni, Transcranial direct current stimulation (tDCS) in the management of epilepsy: A systematic review. Seizure 86, 85–95 (2021).

13. A. R. Brunoni, M. A. Nitsche, N. Bolognini, M. Bikson, T. Wagner, L. Merabet, D. J. Edwards, A. Valero-Cabre, A. Rotenberg, A. Pascual-Leone, R. Ferrucci, A. Priori, P. S. Boggio, F. Fregni, Clinical research with transcranial direct current stimulation (tDCS): Challenges and future directions. Brain Stimulat. 5, 175–195 (2012).

14. S. VanHaerents, B. S. Chang, A. Rotenberg, A. Pascual-Leone, M. M. Shafi, Noninvasive Brain Stimulation in Epilepsy. J. Clin. Neurophysiol. Off. Publ. Am. Electroencephalogr. Soc. 37, 118–130 (2020).

15. N. Grossman, D. Bono, N. Dedic, S. B. Kodandaramaiah, A. Rudenko, H.-J. Suk, A. M. Cassara, E. Neufeld, N. Kuster, L.-H. Tsai, A. Pascual-Leone, E. S. Boyden, Noninvasive Deep Brain Stimulation via Temporally Interfering Electric Fields. Cell 169, 1029–1041.e16 (2017).

16. E. Ben-Menachem, Vagus-nerve stimulation for the treatment of epilepsy. Lancet Neurol. 1, 477–482 (2002).

17. P. Davis, J. Gaitanis, Neuromodulation for the Treatment of Epilepsy: A Review of Current Approaches and Future Directions. Clin. Ther. 42, 1140–1154 (2020).

18. A. Schulze-Bonhage, Brain stimulation as a neuromodulatory epilepsy therapy. Seizure 44, 169–175 (2017).

19. W. H. Theodore, R. S. Fisher, Brain stimulation for epilepsy. Lancet Neurol. 3, 111–118 (2004).

20. V. Salanova, T. Witt, R. Worth, T. R. Henry, R. E. Gross, J. M. Nazzaro, D. Labar, M. R. Sperling, A. Sharan, E. Sandok, A. Handforth, J. M. Stern, S. Chung, J. M. Henderson, J. French, G. Baltuch, W. E. Rosenfeld, P. Garcia, N. M. Barbaro, N. B. Fountain, W. J. Elias, R. R. Goodman, J. R. Pollard, A. I. Tröster, C. P. Irwin, K. Lambrecht, N. Graves, R. Fisher, Long-term efficacy and safety of thalamic stimulation for drug-resistant partial epilepsy. Neurology 84, 1017–1025 (2015).

21. R. Fisher, V. Salanova, T. Witt, R. Worth, T. Henry, R. Gross, K. Oommen, I. Osorio, J. Nazzaro, D. Labar, M. Kaplitt, M. Sperling, E. Sandok, J. Neal, A. Handforth, J. Stern, A. DeSalles, S. Chung, A. Shetter, D. Bergen, R. Bakay, J. Henderson, J. French, G. Baltuch, W. Rosenfeld, A. Youkilis, W. Marks, P. Garcia, N. Barbaro, N. Fountain, C. Bazil, R. Goodman, G. McKhann, K. B. Krishnamurthy, S. Papavassiliou, C. Epstein, J. Pollard, L. Tonder, J. Grebin, R. Coffey, N. Graves, Electrical stimulation of the anterior nucleus of thalamus for treatment of refractory epilepsy. Epilepsia 51, 899–908 (2010).

22. A. L. Benabid, A. Koudsié, A. Benazzouz, L. Vercueil, V. Fraix, S. Chabardes, J. F. LeBas, P. Pollak, Deep brain stimulation of the corpus luysi (subthalamic nucleus) and other targets in Parkinson’s disease. Extension to new indications such as dystonia and epilepsy. J. Neurol. 248, 37–47 (2001).

23. A. L. Benabid, P. Pollak, E. Seigneuret, D. Hoffmann, E. Gay, J. Perret, in Advances in Stereotactic and Functional Neurosurgery 10, Acta Neurochirurgica. B. A. Meyerson, G. Broggi, J. Martin-Rodriguez, C. Ostertag, M. Sindou, Eds. (Springer, Vienna, 1993), pp. 39–44.

24. V. Krishna, N. K. K. King, F. Sammartino, I. Strauss, D. M. Andrade, R. A. Wennberg, A. M. Lozano, Anterior Nucleus Deep Brain Stimulation for Refractory Epilepsy. Neurosurgery 78, 802–811 (2016).

25. F. Velasco, A. L. Velasco, M. Velasco, F. Jiménez, J. D. Carrillo-Ruiz, G. Castro, Deep brain stimulation for treatment of the epilepsies: the centromedian thalamic target. Acta Neurochir. Suppl. 97, 337–342 (2007).

26. A. Cukiert, C. M. Cukiert, J. A. Burattini, A. M. Lima, Seizure outcome after hippocampal deep brain stimulation in a prospective cohort of patients with refractory temporal lobe epilepsy. Seizure - Eur. J. Epilepsy 23, 6–9 (2014).

27. K. Vonck, M. Sprengers, E. Carrette, I. Dauwe, M. Miatton, A. Meurs, L. Goossens, V. De Herdt, R. Achten, E. Thiery, R. Raedt, D. Van Roost, P. Boon, A decade of experience with deep brain stimulation for patients with refractory medial temporal lobe epilepsy. Int. J. Neural Syst. 23, 1250034 (2013).

28. A. Cukiert, C. M. Cukiert, J. A. Burattini, P. P. Mariani, D. F. Bezerra, Seizure outcome after hippocampal deep brain stimulation in patients with refractory temporal lobe epilepsy: A prospective, controlled, randomized, double-blind study. Epilepsia 58, 1728–1733 (2017).

29. F. Velasco, M. Velasco, A. L. Velasco, D. Menez, L. Rocha, Electrical Stimulation for Epilepsy: Stimulation of Hippocampal Foci. Stereotact. Funct. Neurosurg. 77, 223–227 (2001).

30. T. Wyckhuys, T. D. Smedt, P. Claeys, R. Raedt, L. Waterschoot, K. Vonck, C. V. den Broecke, C. Mabilde, L. Leybaert, W. Wadman, P. Boon, High Frequency Deep Brain Stimulation in the Hippocampus Modifies Seizure Characteristics in Kindled Rats. Epilepsia 48, 1543–1550 (2007).

31. T. Wyckhuys, R. Raedt, K. Vonck, W. Wadman, P. Boon, Comparison of hippocampal Deep Brain Stimulation with high (130Hz) and low frequency (5Hz) on afterdischarges in kindled rats. Epilepsy Res. 88, 239–246 (2010).

32. T. Akman, H. Erken, G. Acar, E. Bolat, Z. Kizilay, F. Acar, O. Genc, Effects of the hippocampal deep brain stimulation on cortical epileptic discharges in penicillin - induced epilepsy model in rats. Turk. Neurosurg. 21, 1–5 (2011).

33. P. Boon, K. Vonck, V. D. Herdt, A. V. Dycke, M. Goethals, L. Goossens, M. V. Zandijcke, T. D. Smedt, I. Dewaele, R. Achten, W. Wadman, F. Dewaele, J. Caemaert, D. V. Roost, Deep Brain Stimulation in Patients with Refractory Temporal Lobe Epilepsy. Epilepsia 48, 1551–1560 (2007).

34. N. Axmacher, C. E. Elger, J. Fell, Ripples in the medial temporal lobe are relevant for human memory consolidation. Brain 131, 1806–1817 (2008).

35. M. Mölle, O. Eschenko, S. Gais, S. J. Sara, J. Born, The influence of learning on sleep slow oscillations and associated spindles and ripples in humans and rats. Eur. J. Neurosci. 29, 1071–1081 (2009).

36. J. M. Ibarz, G. Foffani, E. Cid, M. Inostroza, L. Menendez de la Prida, Emergent Dynamics of Fast Ripples in the Epileptic Hippocampus. J. Neurosci. 30, 16249–16261 (2010).

37. A. Bragin, I. Mody, C. L. Wilson, J. Engel, Local generation of fast ripples in epileptic brain. J. Neurosci. Off. J. Soc. Neurosci. 22, 2012–2021 (2002).

38. S. Wang, I. Z. Wang, J. C. Bulacio, J. C. Mosher, J. Gonzalez-Martinez, A. V. Alexopoulos, I. M. Najm, N. K. So, Ripple classification helps to localize the seizure-onset zone in neocortical epilepsy. Epilepsia 54, 370–376 (2013).

39. P. Jiruska, G. T. Finnerty, A. D. Powell, N. Lofti, R. Cmejla, J. G. R. Jefferys, Epileptic high-frequency network activity in a model of non-lesional temporal lobe epilepsy. Brain 133, 1380–1390 (2010).

40. T. Akiyama, B. McCoy, C. Y. Go, A. Ochi, I. M. Elliott, M. Akiyama, E. J. Donner, S. K. Weiss, O. C. Snead, J. T. Rutka, J. M. Drake, H. Otsubo, Focal resection of fast ripples on extraoperative intracranial EEG improves seizure outcome in pediatric epilepsy. Epilepsia 52, 1802–1811 (2011).

41. M. D. Mălîia, C. Donos, A. Barborica, I. Mindruta, I. Popa, M. Ene, S. Beniczky, High frequency spectral changes induced by single-pulse electric stimulation: Comparison between physiologic and pathologic networks. Clin. Neurophysiol. 128, 1053–1060 (2017).

42. M. Lévesque, L. Chen, G. Etter, Z. Shiri, S. Wang, S. Williams, M. Avoli, Paradoxical effects of optogenetic stimulation in mesial temporal lobe epilepsy. Ann. Neurol. 86, 714–728 (2019).

43. ISO/TS 10974:2018(en), Assessment of the safety of magnetic resonance imaging for patients with an active implantable medical device (available at https://www.iso.org/obp/ui/#iso:std:iso:ts:10974:ed-2:v1:en).

44. I. Liorni, E. Neufeld, S. Kühn, M. Murbach, E. Zastrow, W. Kainz, N. Kuster, Novel mechanistic model and computational approximation for electromagnetic safety evaluations of electrically short implants. Phys. Med. Biol. 63, 225015 (2018).

45. F. Missey, E. Rusina, E. Acerbo, B. Botzanowski, A. Trébuchon, F. Bartolomei, V. Jirsa, R. Carron, A. Williamson, Orientation of Temporal Interference for Non-invasive Deep Brain Stimulation in Epilepsy. Front. Neurosci. 15, 633988 (2021).

46. G. Buzsáki, Hippocampal sharp wave-ripple: A cognitive biomarker for episodic memory and planning. Hippocampus 25, 1073–1188 (2015).

47. C. J. Behrens, L. P. van den Boom, L. de Hoz, A. Friedman, U. Heinemann, Induction of sharp wave–ripple complexes in vitro and reorganization of hippocampal networks. Nat. Neurosci. 8, 1560–1567 (2005).

48. R. J. Staba, A. Bragin, High-frequency oscillations and other electrophysiological biomarkers of epilepsy: underlying mechanisms. Biomark. Med. 5, 545–556 (2011).

49. A. E. Musto, M. S. Samii, J. F. Hayes, Different phases of afterdischarge during rapid kindling procedure in mice. Epilepsy Res. 85, 199–205 (2009).

50. C. Huneau, P. Benquet, G. Dieuset, A. Biraben, B. Martin, F. Wendling, Shape features of epileptic spikes are a marker of epileptogenesis in mice. Epilepsia 54, 2219–2227 (2013).

51. S. Demont-Guignard, P. Benquet, U. Gerber, A. Biraben, B. Martin, F. Wendling, Distinct hyperexcitability mechanisms underlie fast ripples and epileptic spikes. Ann. Neurol. 71, 342–352 (2012).

52. N. Roehri, F. Pizzo, S. Lagarde, I. Lambert, A. Nica, A. McGonigal, B. Giusiano, F. Bartolomei, C.-G. Bénar, High-frequency oscillations are not better biomarkers of epileptogenic tissues than spikes. Ann. Neurol. 83, 84–97 (2018).

53. K. Vonck, P. Boon, E. Achten, J. De Reuck, J. Caemaert, Long-term amygdalohippocampal stimulation for refractory temporal lobe epilepsy. Ann. Neurol. 52, 556–565 (2002).

54. D. San-Juan, D. A. Espinoza López, R. Vázquez Gregorio, C. Trenado, M. Fernández-González Aragón, L. Morales-Quezada, A. Hernandez Ruiz, F. Hernandez-González, A. Alcaraz-Guzmán, D. J. Anschel, F. Fregni, Transcranial Direct Current Stimulation in Mesial Temporal Lobe Epilepsy and Hippocampal Sclerosis. Brain Stimulat. 10, 28–35 (2017).

55. H. Jiang, S. Liu, X. Geng, A. Caccavano, K. Conant, S. Vicini, J. Wu, Pacing Hippocampal Sharp-Wave Ripples With Weak Electric Stimulation. Front. Neurosci. 12, 164 (2018).

56. M. I. Iacono, E. Neufeld, E. Akinnagbe, K. Bower, J. Wolf, I. V. Oikonomidis, D. Sharma, B. Lloyd, B. J. Wilm, M. Wyss, K. P. Pruessmann, A. Jakab, N. Makris, E. D. Cohen, N. Kuster, W. Kainz, L. M. Angelone, MIDA: A Multimodal Imaging-Based Detailed Anatomical Model of the Human Head and Neck. PLOS ONE 10, e0124126 (2015).

57. H. McCann, G. Pisano, L. Beltrachini, Variation in Reported Human Head Tissue Electrical Conductivity Values. Brain Topogr. 32, 825–858 (2019).

58. IT’IS Database for Thermal and Electromagnetic Parameters of Biological Tissues – ScienceOpen (available at https://www.scienceopen.com/document?vid=a95fbaa4-efd8-429a-a59e-5e208fea2e45).

59. I. C. on N.-I. R. Protection, ICNIRP Guidelines on Limits of Exposure to Laser Radiation of Wavelengths between 180 nm and 1,000 μm. Health Phys. 105, 271–295 (2013).

60. B. Colombet, M. Woodman, J. M. Badier, C. G. Bénar, AnyWave: a cross-platform and modular software for visualizing and processing electrophysiological signals. J. Neurosci. Methods 242, 118–126 (2015).

61. J. Isnard, D. Taussig, F. Bartolomei, P. Bourdillon, H. Catenoix, F. Chassoux, M. Chipaux, S. Clémenceau, S. Colnat-Coulbois, M. Denuelle, S. Derrey, B. Devaux, G. Dorfmüller, V. Gilard, M. Guenot, A.-S. Job-Chapron, E. Landré, A. Lebas, L. Maillard, A. McGonigal, L. Minotti, A. Montavont, V. Navarro, A. Nica, N. Reyns, J. Scholly, J.-C. Sol, W. Szurhaj, A. Trebuchon, L. Tyvaert, M. P. Valenti-Hirsch, L. Valton, J.-P. Vignal, P. Sauleau, French guidelines on stereoelectroencephalography (SEEG). Neurophysiol. Clin. 48, 5–13 (2018).

62. S. Medina Villalon, R. Paz, N. Roehri, S. Lagarde, F. Pizzo, B. Colombet, F. Bartolomei, R. Carron, C.-G. Bénar, EpiTools, A software suite for presurgical brain mapping in epilepsy: Intracerebral EEG. J. Neurosci. Methods 303, 7–15 (2018).

